# *Beet curly top Iran virus* Rep and V2 gene work as suppressors of post-transcriptional gene silencing through separate mechanisms

**DOI:** 10.1101/2022.08.25.505242

**Authors:** Saeideh Ebrahimi, Omid Eini, Alexandra Baßler, Zeynep Yildirim, Michael Wassenegger, Gabi Krczal, Veli Vural Uslu

## Abstract

Beet curly top Iran Virus (BCTIV) is a yield-limiting geminivirus belonging to the *becurtovirus* genus. The genome organization of BCTIV is unique such that the complementary strand of BCTIV resembles mastreviruses, whereas the virion strand organization is close to curtoviruses. Geminiviruses are known to avoid the plant defense system by suppressing the RNA interference mechanisms both at the transcriptional gene silencing (TGS) and post-transcriptional gene silencing (PTGS) level. Multiple geminivirus genes have been identified as viral suppressors of RNA silencing (VSR) but VSR activity remains elusive in becurtoviruses. By screening all verified open reading frames in the BCTIV genome, we found that only V2 and Rep were able to suppress specific PTGS mechanisms, triggered by the expression of a partial or full-length sense-strand transcript of the target gene (S-PTGS). BCTIV-V2 could suppress S-PTGS more efficiently than BCTIV-Rep when then the target GFP gene is transiently expressed. On the other hand, S-PTGS is suppressed by Rep but not V2 when target GFP is only stably expressed. Deletional mutagenesis of BCTIV-Rep implicated that multiple domains are required for its VSR activity. Furthermore, neither V2 nor Rep could fully suppress local PTGS induced by inverted repeat targeting GFP (GFP-IR). Also, in a closer look at the spread of local silencing by GFP-IR, we observed that V2 or Rep are not able to suppress the movement of sRNAs. Nevertheless, Rep suppressed the systemic silencing induced by GFP-IR in 16C plants. Northern blot analyses showed that BCTIV-Rep inhibits silencing by mitigating sRNA production, whereas BCTIV-V2 does not alter sRNA levels. In summary, both the silencing phenotype and the molecular signatures of silencing implicate distinct modes of VSR activity of BCTIV-Rep and -V2.

## BACKGROUND

*Beet curly top Iran virus* (BCTIV) has a monopartite single-stranded DNA (ssDNA) genome and belongs to *becurtovirus* genus of the *geminiviridae* family (Heydarnejad *et al*. 2013). BCTIV is a yield-limiting virus, causing a serious leaf curl disease in economically important crops such as sugar beet, tomato, pepper, and beans (Yazdi *et al*. 2008, Heydarnejad *et al*. 2013). The leaf curl symptom induced by BCTIV is very similar to those caused by other members of the geminiviridae, such as beet curly top viruses (BCTV) of curtovirus genus (Anabestani et al. 2017). However, BCTIV transmission relies on a distinct vector, *Circulifer haematoceps* (Taheri et al. 2012), and the genome organization of BCTIV differs from the other members of the *geminividae* (Yazdi et al. 2008). Yet the functions of the BCTIV encoded proteins have not been addressed rigorously.

Geminiviruses encode four to eight proteins in their mono- or bi-partite genomes. The transcription from the geminivirus genome takes place in a bidirectional fashion: complementary strand (C) and virion strand (V) (Teixeira et al. 2021). Geminiviruses contain a TAGTATTAC sequence at the replication origin (*ori*), where the replication is initiated by the viral replication initiator protein, known as the complementary strand protein 1 (C1) (Teixeira et al. 2021). Other proteins in the complementary strand are transcriptional activator protein C2, replication enhancer protein C3, and pathogenicity determinant protein C4 (Kumar 2019). In addition, in *mastrevirus, capulavirus*, and *grablovirus* genera, due to a splicing event between C1 and C2, a new open reading frame (ORF), called Rep, emerges containing the N terminus of C1 and a frame-shifted C terminus of C2. Rep protein of geminiviruses is a multifunctional protein, which is indispensable for the viral genome replication (Ruhel and Chakraborty 2019). On the other hand, the virion strand encodes the coat protein (V1), movement protein (V2), and a poorly characterized gene (V3) (Kotlizky et al. 2000). The C and V genes are organized in unique combinations for each genus of the geminiviridae family. For example, BCTIV contains C1, C2, and Rep on the complementary-sense strand, similar to Mastreviruses, whereas V1, V2, and V3 on the virion-sense strand, similar to curtoviruses (Bozorgi et al. 2017, Zerbini et al. 2017) (Supplementary Figure 1a).

Geminivirus pathology is influenced by the efficiency of virus accumulation, which predominantly depends on viral suppression of RNA interference (RNAi) in plants (Hanley-Bowdoin et al. 2013). RNAi is a conserved defense mechanism, acting through transcriptional (TGS) or post-transcriptional (PTGS) repression of gene expression in a sequence-specific manner (Baulcombe 2004). Bi-directional transcription from the geminivirus genome creates double-stranded RNA (Vanitharani et al. 2005). In addition, aberrant geminivirus transcripts can be used as templates for dsRNA production by plant RNA-dependent RNA polymerases (RDRs) (Bisaro 2006). dsRNAs are cleaved into small RNAs (sRNAs) of distinct lengths by Type III endonucleases, called DICER-LIKE (DCL) proteins (Baulcombe 2004). sRNAs are incorporated into ARGONAUTE (AGO) proteins and form the RNA-induced silencing complex (RISC) for the cleavage of the target sequence. Besides cleavage, RISC can also control PTGS by recruiting RDRs for amplification of the RNAi or inhibiting target gene translation facultatively (Yoshikawa et al. 2021, Uslu et al. 2021, Wu et al. 2020). In addition, auxiliary RNAi proteins like HUA ENHANCER 1 (HEN1) RNA methyltransferase, which protects sRNAs from degradation, contribute to viral defense in plants (Boutet et al. 2016, Muangsan et al. 2004, Jin et.al 2022)

There are many identified viral PTGS suppressors in plant viruses. For example, helper component-proteinase (HC-Pro) encoded by *Potyviridae* inhibits RDR6 (Zhang et al. 2008) and HEN1 (Jamous et al. 2011, Sanobar et al. 2021) among other components of the RNAi pathway, whereas, P19 protein (P19) of *Tomato bushy stunt virus* (TBSV) sequesters ds-siRNAs inhibiting RNAi based protection in plants (Voinnet et al. 2003, Ye et al. 2003). In addition, the P6 protein of *Strawberry vein banding virus* (Feng et al. 2018), the P20, P23, and capsid proteins of *Citrus tristeza virus* (Lu et al. 2004), the P6 protein of *Cauliflower mosaic virus* (Haas et al. 2008), P38 protein of *Turnip crinkle virus* (Iki et al. 2017), P1 of *Sweet potato mild mottle virus* (Giner et al. 2010) are among those, which are have been shown to function as viral suppressors of RNA silencing (VSR). In *Geminiviridae*, several reports have implicated that C2, C4, Rep, V2, and V3 proteins (Liu et al. 2014, Wang et al. 2014, Hanley-Bowdoin et al. 2013) function as viral VSRs, involved in the suppression of transcriptional gene silencing (TGS) or post-transcriptional gene silencing (PTGS) in plants. Yet, the silencing suppressor activity of the BCTIV gene products remains elusive.

In this study, we induced PTGS using specific silencing inducers, including full-length GFP, partial sense-strand GFP, and inverted-repeat GFP to silence GFP expression. By combining the given silencing inducers with each of the BCTIV open reading frames on *Nicotiana benthamiana*, we investigated the efficiency of PTGS in local and systemic contexts to find out the BCTIV genes, responsible for PTGS suppression.

## METHODS

### Plasmid constructs for transient expression

Sense-PTGS silencing inducers were cloned into pGreen (pG104) vector containing full length (FL-GFP) or only 139nt long 5′-GFP sequence (5′-GFP) (Dalakouras et al. 2019). Strong PTGS was induced by the inverted repeat of the very same 139nt sequence (GFP-IR) with a 90nt long spacer as described in Dalakouras et al. 2019. The positive VSR control P19 of TBSV was expressed in pPCV702SM binary vector (Saeed et al. 2015). All silencing inducers are expressed under the control of 35S promoter.

For unbiased investigation of the genes potentially responsible for VSR all verified ORFs of BCTIV (C1, C2, C4, Rep, V1, V2, V3) in pDIVA and/or pG104 binary vectors (Ebrahimi et al. 2022) were examined on 16C plants (Supplementary Figure 1). The coding sequences of the C1, C2, Rep, and V2 genes were amplified with primers containing *BamHI* and *XhoI* restriction cut sites (Table 1), using Q5 polymerase (NEB). The PCR products were first cloned into pJET1.2/blunt cloning vector (Thermo Fischer Scientific, Germany) and they were sub-cloned using *Xho*I and *BamHI* sites of pG104 vector to obtain pG-C1, pG-C2, pG-Rep and pG-V2, to obtain the same vector backbone for all silencing inducers. The inserts were confirmed by Sanger sequencing. Of note, we found out that the vector backbone did not influence the silencing or VSR phenotypes.

12 truncations from the Rep gene were PCR-amplified using corresponding primers (Table1) and cloned into pJET1.2. Then, the plasmids with Rep truncations were digested with the *BamHI* and *XhoI* enzymes they were cloned into the pG104 vector backbone to obtain pG-Rep1, pG-Rep2, pG-Rep3, pG-Rep4, pG-Rep5, pG-Rep6, pG-Rep7, pG-Rep8, pG-Rep9, pG-Rep10, pG-Rep11, and pG-Rep12 (Supplementary Figure 2). *Escherichia coli* (strain INVαF, ThermoFisher) was used for cloning plasmid constructs and *A. tumefaciens* strain ATHV was used for transient transformation of the plants. pSoup helper plasmid was used together with pG104 for transforming agrobacterium by electroporation. In addition to the plasmid constructs obtained from INVαF, plasmids in the *agrobacterium* clones were also sequenced to ensure the insert integrity.

### Agroinfiltration

The BCTIV genes were co-infiltrated with *Agrobacterium* ATHV containing silencing inducer constructs into wildtype *Nicotiana benthamiana* (WT) and GFP-expressing *N*.*benthamiana* (16C) at the four/six-leaf-stage. *Agrobacteria* containing the constructs were grown in LB medium (containing Kanamycin and Rifampicin) at 28.0 °C for 16 h with continuous shaking at 135 rpm, then re-suspended in MES buffer (10 mM MgCl2, 10 mM MES (pH 5.6), and 100 μM acetosyringone) to reach OD_600_= 1 for agroinfiltration. For negative control, *N. benthamiana* plants were separately infiltrated with FL-GFP, 5′-GFP, and GFP-IR. In order to equalize the strength of silencing induction in single silencing inducer infiltrations, agrobacterium containing empty pG104 (OD_600_= 1) has been added for mock co-infiltrations. Unless otherwise stated, three plants were used for each treatment in each experiment, and each experiment was repeated at least three times at different times of the year to avoid any effect of temperature fluctuations in the greenhouse.

### GFP imaging and quantification

The inoculated plants were maintained in a growth chamber at 25 °C with 16-h light/ 8-h dark period. After 3 days post inoculation (dpi) the control plants were monitored regularly under UV light (Blak-Ray B-100 AP Lamp, www.uvp.com) to visualize the silencing phenotype. At 6dpi all the plants were recorded using a Canon camera (Canon EOS700D, 18-5mm Lens) under UV light and leaves were scanned using a laser scanner (Molecular Imager® PharosFX™ Systems, BioRad) at 50µm per pixel, in GFP (excitation at 488nm, emission at 530nm) and in Chlorophyll (excitation at 488nm, emission at 695nm) channels.

The GFP and the Chlorophyll images were merged and then GFP expression was quantified in the limited area of infiltrated patches. GFP expression was measured by FIJI image analysis (www.imagej.net) in treated, non-necrotic area and the acquired values were normalized using non treated areas. The final values were plotted in bar charts (BoxPlotR, Shiny Lab).

### Protein Extraction and Western blot

Total protein was extracted from 6 dpi infiltrated leaf patches of WT plants using protein extraction buffer (1M Tris-HCl, PH 8.8, 10% Glycerol, 10% SDS, and H2O) together with cOmplete^™^ Protease inhibitor solution (Roche). SDS-PAGE was performed for Coomassie staining and Western blotting (Wb), polyclonal goat anti-GFP antibody (Sicgen, AB0066-200, 1:1000) as primary, Donkey anti-goat IgG-HRP antibody (sc-2020, Santa Cruz Biotechnology, (1:5000) as secondary antibody were used for western blot. Pierce™ ECL Western Blotting Substrate (ThermoFisher) was used to detect the signal. 16C *N*.*benthamiana* were not included for the western blot analysis due to the fact that the vasculature of 16C is barely affected by silencing and depending on how much vasculature is included in the western blot sample collection from silenced areas, the results fluctuate dramatically.

### RNA extraction, RT-PCR, and Northern blot

Total RNA was extracted from infiltrated patches of 16C leaves at 6 dpi using TRIzol® Reagent (Qiagen). RNA concentration and RNA purity were measured using Nanodrop (ND-1000) and RNA quality was assessed by agarose gel electrophoresis. RNAs are treated with Lucigen Baseline-ZERO DNase and purified by acid-Phenol-Chloroform pH:4.5 (Invitrogen™, Thermofisher). RT-PCR was performed using the SuperScript™ III One-Step RT-PCR System with Platinum™ Taq DNA Polymerase (Invitrogen™, ThermoFisher) and gene-specific primers (Table 2) to amplify BCTIV genes to confirm transient expression of the constructs upon agroinfiltration. For the Northern blot of GFP mRNA and GFP siRNA 10μg and 5μg of total RNA were used, respectively, as previously described (Dalakouras et al. 2019). The siRNA Northern blots were stripped and rehybridized with a U6 snoRNA probe. For GFP probes [alpha-P32] CTP is used for labeling, for U6 snoRNA probe [gamma-p32] ATP is used from Hartman Analytic, Germany. The detection of the Northern blot signals was performed by the Molecular Imager® PharosFX™ Systems (BioRad).

### Statistical Analysis

Statistical tests have been performed by One-way ANOVA using JMP Software (https://www.jmp.com/). Unless otherwise stated, p<0.05 is taken as the significance threshold.

### Phylogenetic Tree

Nucleotide sequences of the Rep (or homologous) proteins of different geminiviruses were obtained from NCBI. MEGA (Molecular Evolutionary Genetics Analysis) was used for the phylogenetic tree, based on the neighbor-joining method.

## RESULTS

### Phenotypic analysis of transient BCTIV gene expression and silencing inducers on *N. benthamiana*

Before we began investigating the VSR activity of BCTIV genes, we tested the expression and the effect of the BCTIV genes upon agroinfiltration into 16C *N. benthamiana* leaves. C1, C2, Rep, V1, V2, and V3 genes, which were previously shown to be expressed by BCTIV, and C4 was shown to be expressed in BCTV. (Supplementary Figure 1a) (Eini et al. 2016, Yazdi et al. 2008, Teng et al. 2010). Using RT-PCR, we confirmed the expression of the viral genes in *N. benthamiana* leaves (Supplementary Figure 1b). Notably, C1 and Rep expression also led to necrotic speckles starting from 5dpi and the necrotic phenotype gradually got stronger. As sense-PTGS (S-PTGS) inducer, the 5’-GFP construct was agroinfiltrated into GFP expressing 16C leaves. The silencing phenotype induced by 5’-GFP was most consistently visible earliest at 6 dpi. Therefore, the effect of BCTIV genes on local PTGS was examined at 6 dpi.

### BCTIV-Rep suppresses local sense-PTGS in *N*.*benthamiana*

First, we aimed at investigating the role of BCTIV genes in the Sense-PTGS (S-PTGS) pathway. For this assay, 139nt long sequence mapping to GFP (5’-GFP) was transiently expressed together with the BCTIV genes in 16C *N. benthamiana*. Due to S-PTGS induction, agroinfiltrated leaf sectors turn red under UV light due to loss of GFP at 6 dpi. However, if a BCTIV gene functions as a viral suppressor of RNA silencing (VSR) the agroinfiltrated sector should remain green under UV light (Figure 1a). Based on the GFP expression level under the UV light at 6 dpi, we found out that only Rep protein could suppress the S-PTGS, while none of the other constructs, including the positive control TBSV-P19, could suppress S-PTGS (Figure 1b). For the exact quantification of GFP levels after silencing induction and silencing suppression, the leaves were imaged in a high-resolution laser scanner (Figure 1c). The GFP intensity measurements in exclusively non-necrotic areas showed that GFP expression was significantly (*P* < 0.05) higher in Rep infiltrated 16C *N. benthamiana* leaves in comparison to TBSV-P19 and other BCTIV gene infiltration plants. (Figure 1d).

**Figure 1.**
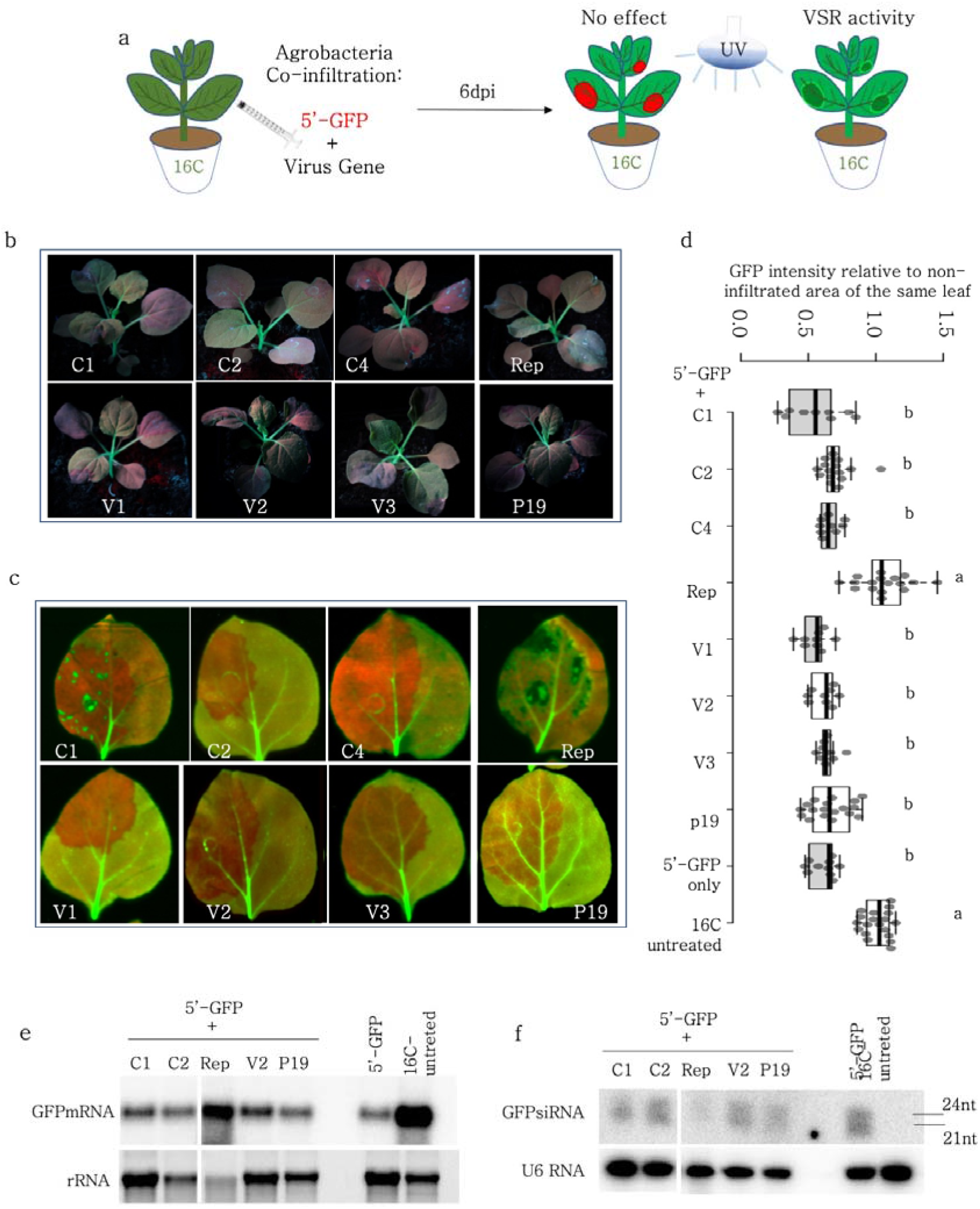
Screening of BCTIV genes for VSR activity. a) GFP expressing 16C *N*.*benthamiana* leaves were co-infiltrated with S-PTGS inducer 5’-GFP and a single BCTIV gene for each experiment. At 6dpi, if 5’-GFP leads to red sectors under the UV lamp, it suggests that the VSR activity of the co-infiltrated gene is negligible. However, if the infiltrated sector remains green under UV lamp, it shows VSR activity for the gene of interest. b) Whole plant photos of the co-infiltrated plants were taken under UV light by a DLSR camera at 6dpi. Representative images of at least three biological replicates are provided. c) High-resolution images of the leaves infiltrated by 5’-GFP and C1, C2, C4, Rep, V1, V2, and V3 are obtained at 50µm/pixel using green and far-red channels of PharosFX™ imager. Green (GFP) and far-red channels (chlorophyll) are merged d) Non-vasculature sectors of the high-resolution leaf images are quantified based on GFP. At least three leaves and multiple areas per leaf were quantified. One-way ANOVA statistical test shows significance at p<0.05 e) Top panel shows northern blot results using a radioactively labelled GFP probe. 5’-GFP serves as a low GFP expression control and a 16C-untreated sample shows high-abundance GFP mRNA control. rRNA is a loading control for the upper panel, obtained by agarose gel electrophoresis using ethidium bromide staining. f) Top panel shows small RNA Northern blot using a radioactively labelled GFP probe. The bands correspond to 21 and 24 nt long siRNAs. 5’-GFP serves as a highly abundant siRNA control and 16C-untreated serves as a clean negative control. The lower panel is the small RNA Northern blot by using U6 RNA labeling as a control for RNA loading. U6 RNA labelling took place on the same membrane given in the panel above, after stripping.

To reveal the mode of action of VSR activity of BCTIV-Rep, we investigated the GFP mRNA and small RNA populations upon co-infiltration in the leaf sections of 16C plants using Northern blot. To compare with Rep, we used a restricted set of BCTIV-genes, including C1 and C2 because of overlapping sequences with Rep, and V2 genes due to the VSR activity of V2 homologs in other geminiviruses (Luna et al. 2017, Zhang et al. 2012, Zrachya et al. 2006). Despite loading the same amounts of material according to nanodrop measurements, the RNA quality of different co-infiltrations differed observably based on rRNA bands on the agarose gel electrophoresis of the RNA samples. (Figure 1e). Especially, due to the necrotic areas caused by Rep, the RNA integrity was the lowest in Rep + 5′-GFP co-infiltrated sample. Nevertheless, the GFP mRNA level in Rep + 5′-GFP was the highest when compared to other co-infiltrated samples (Figure 1e).

In addition, we performed sRNA Northern Blot on the very same co-infiltrated samples. End labeled U6 snoRNA probe was used as a loading control, which did not reveal any major fluctuations of sRNA concentration (Figure 1f). Samples from untreated 16C did not show any sRNAs, while 5′-GFP treated 16C samples had a clear accumulation of sRNAs, suggesting that the experimental setup was valid. Rep + 5′-GFP sample showed a very faint sRNA band at the level of 21-24 nucleotides (nt), suggesting that Rep interfered with the silencing pathway, before the production of sRNAs by DCLs (Figure 1f). P19 and V2 co-infiltration with 5′-GFP led to only a limited decrease in sRNA production in concordance with the previous literature (Saeed et al. 2015).

### BCTIV-V2 suppresses local S-PTGS only when the target GFP is transiently expressed

In order to detect the changes in GFP expression and silencing levels in a clean background with no GFP fluorescence over the whole leaf surface, full-length GFP (GFP-FL) was co-infiltrated together with BCTIV-genes in WT *N. benthamiana*. GFP-FL under the control of the 35S promoter induces GFP expression. However, due to the S-PTGS mediated self-silencing of the transient GFP-FL expression, the GFP signal remains low. When a BCTIV gene increases the GFP expression, upon co-infiltration with GFP-IR, this should be due to the VSR activity of the gene, which inhibits S-PTGS, hence, the self-silencing of GFP-FL (Figure 2a).

**Figure 2.**
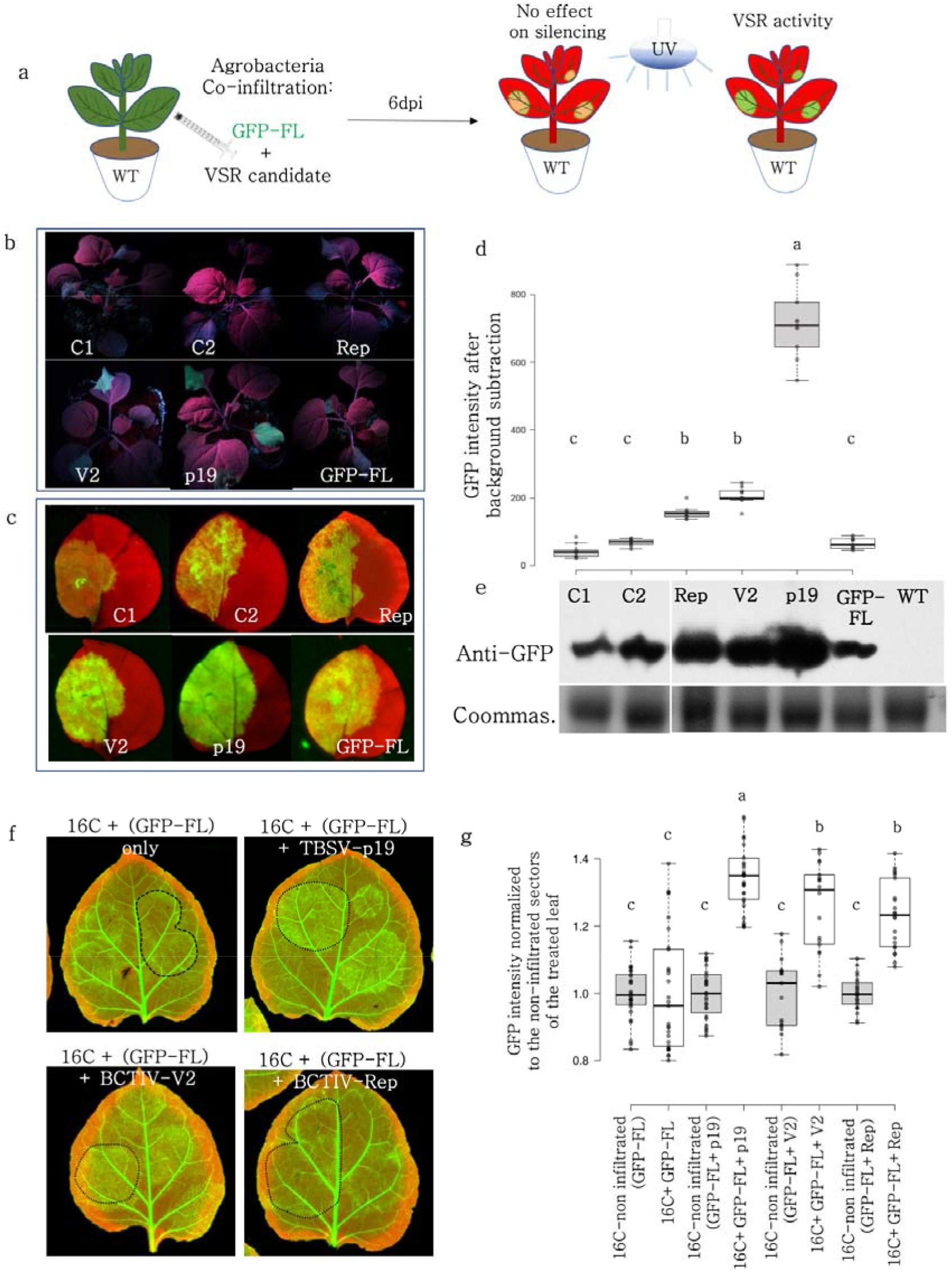
BCTIV-Rep and BCTIV-V2 genes repress S-PTGS upon transient expression of GFP. a) S-PTGS inducing FL-GFP is co-infiltrated with C1, C2, Rep, and V2 of BCTIV on WT *N*.*benthamiana*. Upon VSR activity, GFP-FL agroinfiltration is expected to show high GFP fluorescence under UV light. b) Whole plant photos of the co-infiltrated plants taken by a DLSR camera at 6dpi are shown. TBSV-P19 co-infiltration with FL-GFP is used as a positive control for very high GFP expression and FL-GFP single infiltration serves as a low GFP expression control. c) High-resolution images of the leaves infiltrated by GFP-FL and virus genes are obtained at 50µm/pixel using green and far-red channels of PharosFX™ imager. Green (GFP) and far-red channels (chlorophyll) are merged using FIJI. d) Infiltrated areas in the leaves are quantified based on GFP intensity. At least three leaves and multiple areas per leaf were quantified. One-way ANOVA statistical test shows significance at p<0.05. e) The top panel is the Western blot result, showing the abundance of GFP protein in the co-infiltrated sections of the very same leaves shown in c. the lower panel is the Rubisco band in the Coomassie staining as a loading control for the Western blot. Since the assay is performed on WT *N*.*benthamiana*, it allows quantification of absolute changes in GFP level, and therefore a reliable comparison of image quantifications and protein abundance in Western blot f) FL-GFP is co-infiltrated with p19, V2, and Rep. Representative images from at least two plants and multiple leaves are provided. g) The GFP level quantification of the co-infiltrated sectors normalized to the non-infiltrated sectors of the treated leaves. One-Way ANOVA test has been conducted with a threshold of p<0.05.

Plants under the UV light at 6 dpi showed high GFP expression in the BCTIV-V2 and BCTIV-Rep treated plants compared to only GFP-FL infiltrated plants (Figure 2b). The strongest green fluorescence was observed in TBSV-P19 treated plants, in accordance with the previous literature (Figure 2b) (Ahn et al. 2011, Saeed et al. 2015, Zhang 2015). The quantification based on high-resolution images of infiltrated leaves also corroborated with the observations under the UV lamp, in which P19, V2, and Rep showed higher GFP signal than the GFP-FL only and GFP-FL with C1 or C2 (Figure 2c, d).

Since WT plants provided a blank background for GFP analysis, we addressed whether the quantifications on the images based on GFP intensity in non-necrotic areas matched with the GFP amount in the non-necrotic areas obtained by the Western blot of the infiltrated areas. The Western Blot results showed that co-infiltration of GFP-FL with P19, V2, and Rep led to higher GFP protein expression than GFP-FL alone at 6dpi, in the same order of intensity (Figure 2e). Moreover, C1 and C2 co-infiltration with GFP-FL did not increase the GFP protein level in Western blot in accordance with the image analysis.

We performed the GFP-FL co-infiltration with P19, V2, and Rep on GFP expressing 16C *N*.*benthamiana*. GFP-FL increases GFP expression but also functions as an S-PTGS inducer in the later stages. At 5.25dpi, we found out that the net contribution of GFP-FL on the GFP level is zero (Figure 2f, g). At this stage, P19 works as a silencing suppressor, more efficiently than V2. Although V2+GFP-FL tends to have a stronger silencing suppressor effect than Rep+GFP-FL on average, the difference is not statistically significant (Figure 2g).

### Neither Rep nor V2 can suppress PTGS induced by a hairpin construct targeting GFP

Since Rep is capable of decreasing sRNA production, necessary for S-PTGS, we addressed whether Rep could repress PTGS induced by hairpin RNA (GFP-IR) expressed under the control of the 35S promoter. Similar to the S-PTGS infiltrations using GFP-FL and 5′GFP, the same agrobacterium batch containing BCTIV genes (C1, C2, Rep and V2) were co-infiltrated with GFP-IR in stably GFP expressing 16C plants (Figure 3a). GFP-IR is a very strong inducer of silencing and at 6dpi, none of C1-, C2-, Rep-, V2-, P19-co-infiltrated leaves showed loss of the silencing phenotype (Figure 3b). Nevertheless, C1 and Rep co-infiltrated with GFP-IR had a slightly higher level of GFP than GFP-IR only (Figure 3c, d). Therefore, we investigated the molecular fingerprints of BCTIV genes on PTGS by checking GFP mRNA and sRNA levels using Northern blot on stably GFP expressing 16C plants. Due to lower necrotic areas in Rep + GFP-IR sample, the RNA quality was comparable to the other treatments and controls. In addition, the abundance of GFP mRNA in 16C and the low level of GFP in GFP-IR treatment validated the experimental set-up. None of the treatments, including Rep, could block the decrease of GFP mRNA level upon GFP-IR induction (Figure 3e).

**Figure 3.**
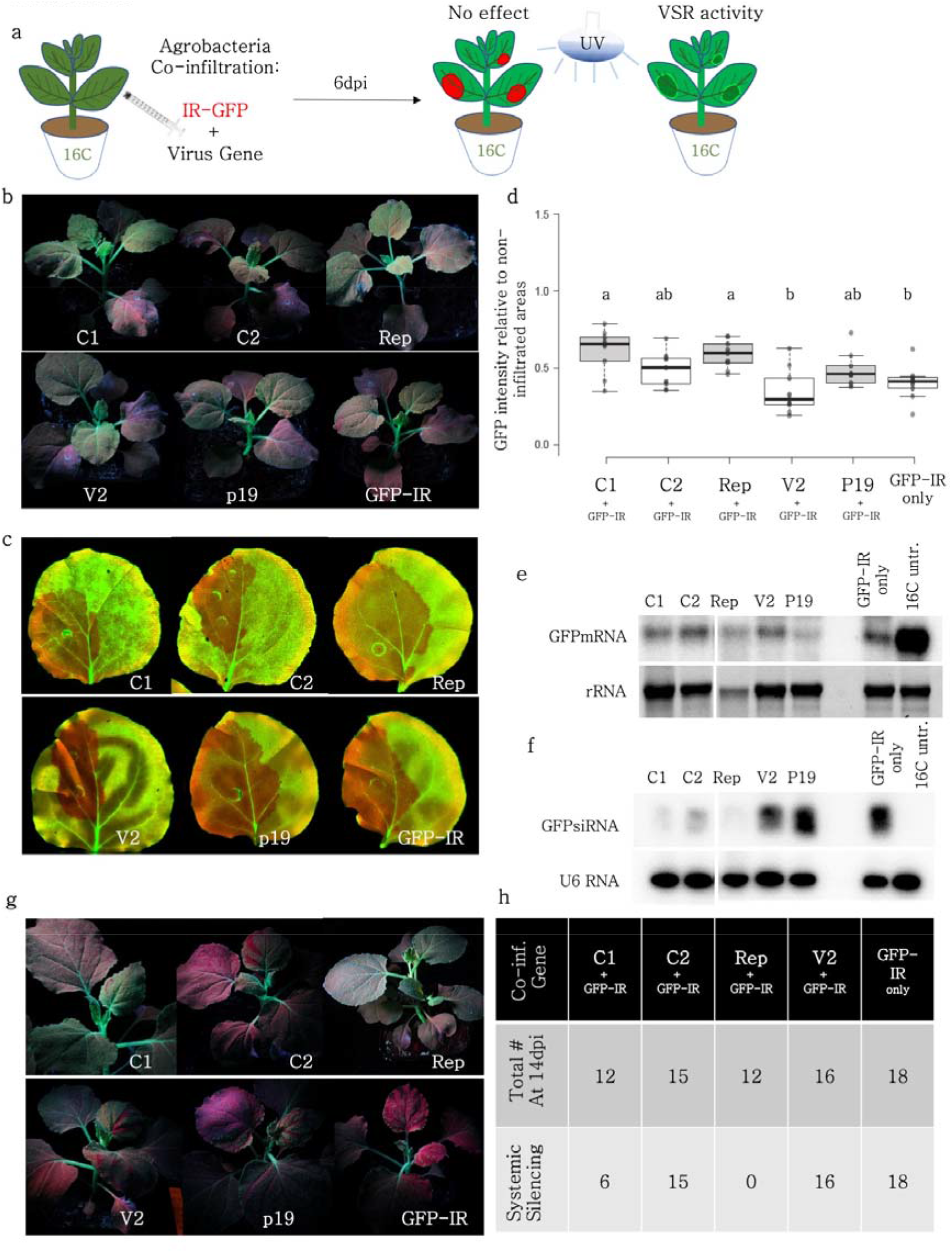
Neither BCTIV-Rep nor BCTIV-V2 can suppress PTGS induced by GFP-IR. a) PTGS-inducing GFP-IR is co-infiltrated with C1, C2, Rep, and V2 of BCTIV on 16C *N*.*benthamiana*. Upon VSR activity, GFP-IR agroinfiltrated sectors are expected to retain high GFP fluorescence under UV light. b) Whole plant photos of the co-infiltrated plants taken under UV light by a DLSR camera at 6dpi are shown. c) High-resolution images of the leaves infiltrated by GFP-IR and C1, C2, Rep, V2, and p19 are obtained at 50µm/pixel using green and far-red channels d) Non-vasculature sectors of the high-resolution leaf images are quantified based on GFP intensity. At least three leaves and multiple areas per leaf were quantified. e) Top panel shows northern blot results using a radioactively labelled GFP probe. GFP-IR serves as a low GFP expression control and a 16C-untreated sample shows high-abundance GFP mRNA control. rRNA is a loading control for the upper panel, obtained by agarose gel electrophoresis using ethidium bromide staining. f) Top panel shows small-RNA-Northern blot using a radioactively labelled GFP probe. The bands correspond to 21 and 24 nt long siRNAs. GFP-IR serves as a highly abundant siRNA control and 16C-untreated serves as a negative control. The lower panel is the small RNA Northern blot by using U6 RNA labelling as a control for RNA loading. U6 RNA labelling took place on the same membrane given in the panel above, after stripping. g) A minimum of twelve 16C *N*.*benthamiana* used for GFP-IR co-infiltration with VSR candidates are followed for until 14 dpi. The images of the whole plant are taken using a DSLR Camera under UV h) The total number of co-infiltrated plants, remaining healthy at 14dpi is provided on the top row, and the number of plants showing systemic silencing in at least two leaves is given the bottom row.

Next, we performed an sRNA Northern blot on GFP-IR induced PTGS samples. U6 snoRNA loading control shows comparable amounts of total sRNAs among different treatments and controls (Figure 3f). In addition, GFP-IR only shows very strong sRNA accumulation, whereas no sRNAs are visible in 16C control, validating the experimental set-up. Interestingly, we have seen that the sRNAs mapping to GFP are almost absent in Rep + GFP-IR and dramatically reduced in C1+GFP-IR at 6dpi. On the other hand, V2 and P19 co-expression with GFP-IR did not have any effect on sRNA accumulation at 6dpi (Figure 3f).

### can suppress systemic silencing 2Rep but not V

To find out whether the decrease in sRNA amounts upon expression of BCTIV genes suppresses systemic silencing in 16C plants (Figure 3f), the GFP-IR treated plants were observed and recorded regularly under UV light until 14dpi (Figure 3g). Systemic silencing emerged at 10 dpi in plants infiltrated with GFP-IR only, upon C2+GFP-IR and V2+GFP-IR co-infiltration the systemic silencing phenotype spread in the newly emerging leaves at 14 dpi, without any exception (Figure 3g, h). However, none of the eighteen plants co-infiltrated with Rep+GFP-IR showed systemic silencing (Figure 3h). These results indicate Rep protein suppression activity in systemic PTGS. It is noteworthy that twelve out of eighteen C1+GFP-IR treated plants did not show any systemic silencing and the remaining six plants showed rather restricted systemic spread of the silencing signal (Figure 3g, h).

### Rep and V2 do not interfere with the spread of silencing through plasmodesmata

Local silencing induced by GFP-IR and/or GFP-FL creates a visible area encircling the infiltration zone, which shows stronger silencing than the infiltration zone (Figure 4a, Himber et al. 2003). It is postulated that the silencing signal reaches this area via the short-range spreading of sRNAs through plasmodesmata (Dunoyer et al. 2005, Kalantidis et al. 2008). It is possible to quantify the length and GFP intensity (Gray value) of this area by plotting the profile (using FIJI) along a line in the GFP channel. Our data showed that the GFP-IR+GFP-FL co-infiltration created a silenced zone of 0.7 mm around the infiltration zone, which roughly corresponds to 15 cells (Figure 4b). When GFP-FL, GFP-IR are co-infiltrated with p19, V2 or Rep on 16C *N*.*benthamiana*, the silencing phenotype is partially repressed at 5.25dpi (Figure 4c, e, g, i). P19 abolishes the silenced ring around the infiltrated area (Figure 4d). However, the silenced area is preserved when BCTIV-V2 and BCTIV-Rep are co-infiltrated with the GFP-IR and GFP-FL (Figure 4f, g).

**Figure 4.**
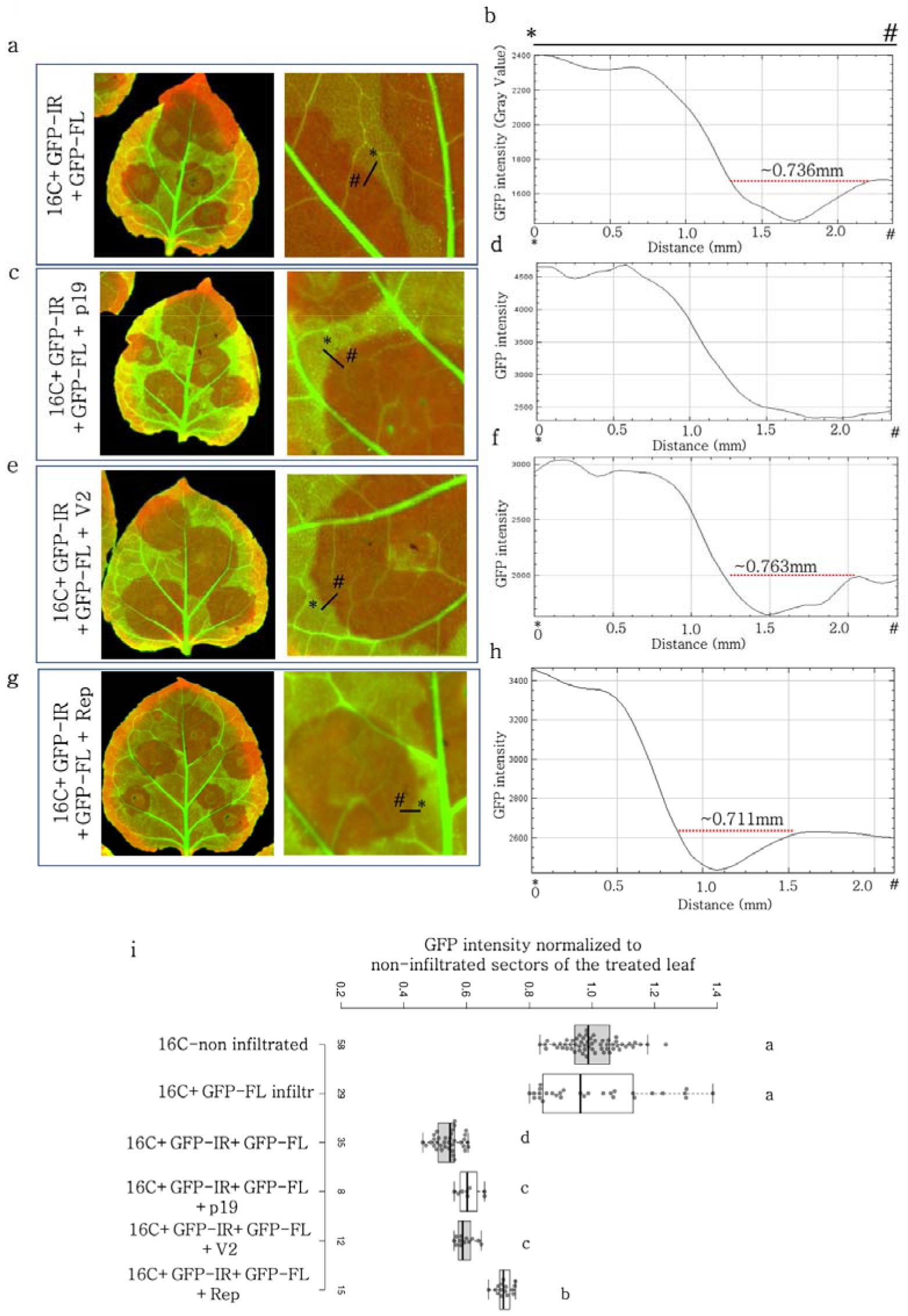
BCTIV-Rep or -V2 does not block the short-range spreading of silencing upon IR induced PTGS. a-g) 16C plants are infiltrated with GFP-FL and GFP-IR only(a), GFP-FL, GFP-IR, and TBSV-p19(b), GFP-FL, GFP-IR, and BCTIV-V2(c), GFP-FL, GFP-IR, and BCTIV-Rep(d). The images are overlay of green and far-red channels, acquired by PharosFX™ imager at 5.25dpi. The close-up on the right highlights the red sector around the infiltration area. b-h) GFP intensity profile of a 10pixel-wide line depicted on the close-up image on the right. Y-axis gives the intensity of the GFP in gray value, X-axis is the distance (in mm) of the spot to the * end of the line given. The left end of the x-axis is the # end of the line. The red dotted line indicates the distance of the short-term spread of local silencing in mm. i) The GFP intensity measurement of infiltrated non-vasculature areas is normalized to their non-infiltrated areas in the same leaf. 16C + GFP-FL infiltrated measurements are taken from Figure 1g for comparison. The corresponding measurement number is given below the bar. One-Way ANOVA test has been conducted with a threshold of p<0.05.

### Multiple Rep domains are indispensable for VSR activity

To figure out if there is any specific domain responsible for suppression activity in Rep protein, different truncations of the Rep gene were constructed based on C1 and frameshift C2 motifs (Supplementary Fig 2). The Rep truncations were co-expressed together with 5′-GFP in 16C *N. benthamiana* plants (Figure 5a). None of the 12 Rep truncations recapitulated the Rep-VSR activity at 6 dpi (Figure 5b) which correlates with GFP quantification in infiltrated leaf patches (Figure 5 c,d). Noteworthy that the 30 amino acid long deletion at the C-terminus of Rep partially kept the VSR activity, suggesting that the C terminus could have minor functions for S-PTGS suppression. Besides, N-terminus deletions but not C-terminus deletions of Rep truncations showed necrotic phenotypes comparable to Full-length Rep protein. These results suggest that multiple domains BCTIV-Rep domains except the very end of C-terminus, are indispensable for silencing function, but the N-terminus of the Rep protein is essential for necrosis, which is similar to wheat dwarf virus (WDV) Rep protein (Liu et al. 2014)

**Figure 5.**
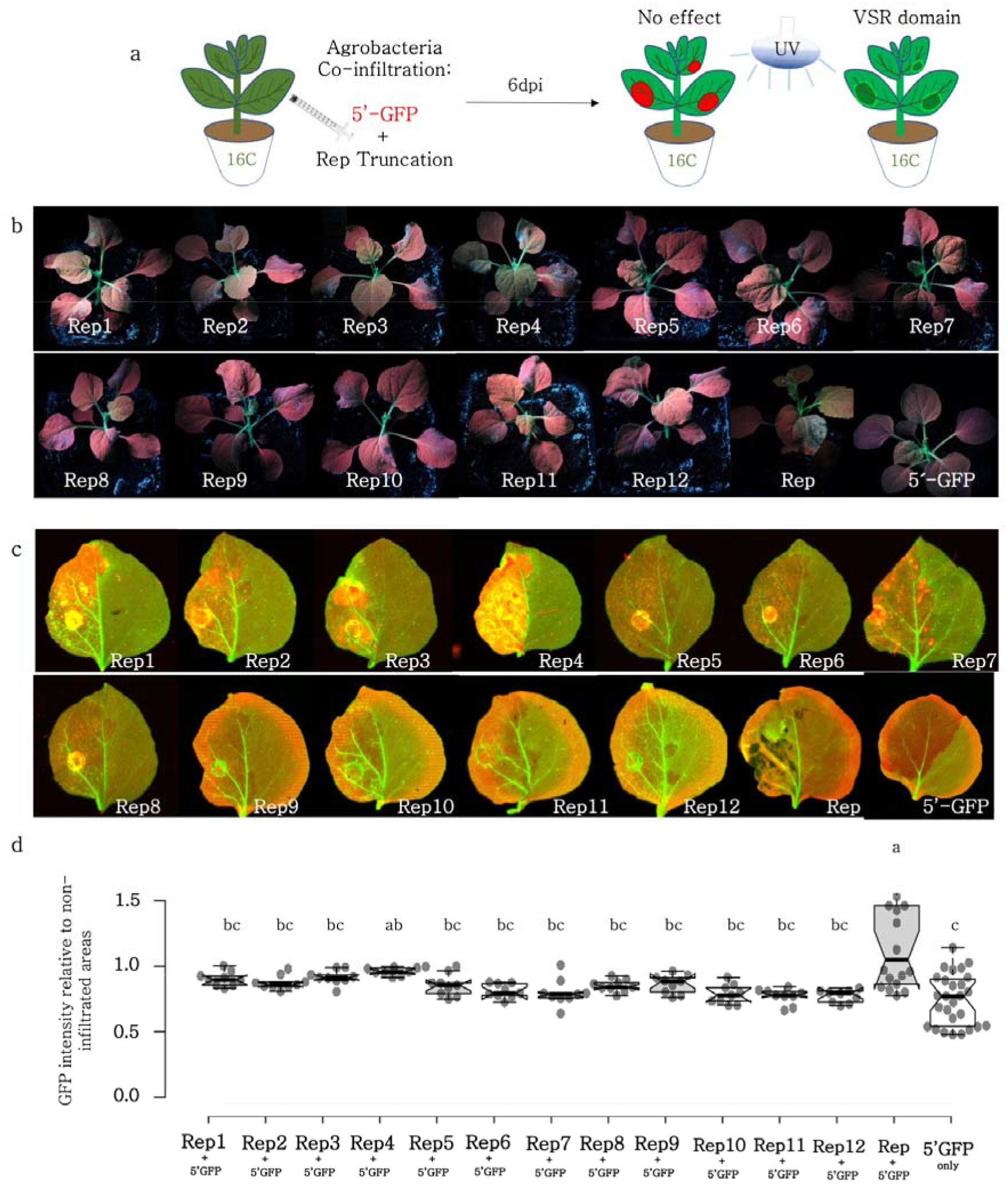
Multiple domains are needed for the VSR activity of BCTIV-Rep. a) GFP expressing 16C *N*.*benthamiana* leaves were co-infiltrated with S-PTGS inducer 5’-GFP and a single BCTIV-Rep truncation in each experiment. At 6dpi, if 5’-GFP leads to red sectors under the UV lamp, it suggests that the VSR activity of the co-infiltrated gene is negligible. However, if the infiltrated sector remains green under UV lamp, it shows VSR activity for the gene of interest. b) Whole plant photos of the co-infiltrated plants were taken under UV light by a DLSR camera at 6dpi. Representative images of the two biological replicates are provided for each co-infiltration. c) High-resolution images of the co-infiltrated leaves are obtained at 50µm/pixel using green and far-red channels of PharosFX™ imager. Green (GFP) and far-red channels (chlorophyll) are merged d) The GFP intensity of the non-vasculature sectors of the high-resolution leaf images are normalized to the non-infiltrated side of the same leaf. At least two leaves and multiple areas per leaf were quantified. The values are plotted by the online BoxPlotR tool by Shiny Lab. One-way ANOVA statistical test shows significance at p<0.05.

**Figure 6.**
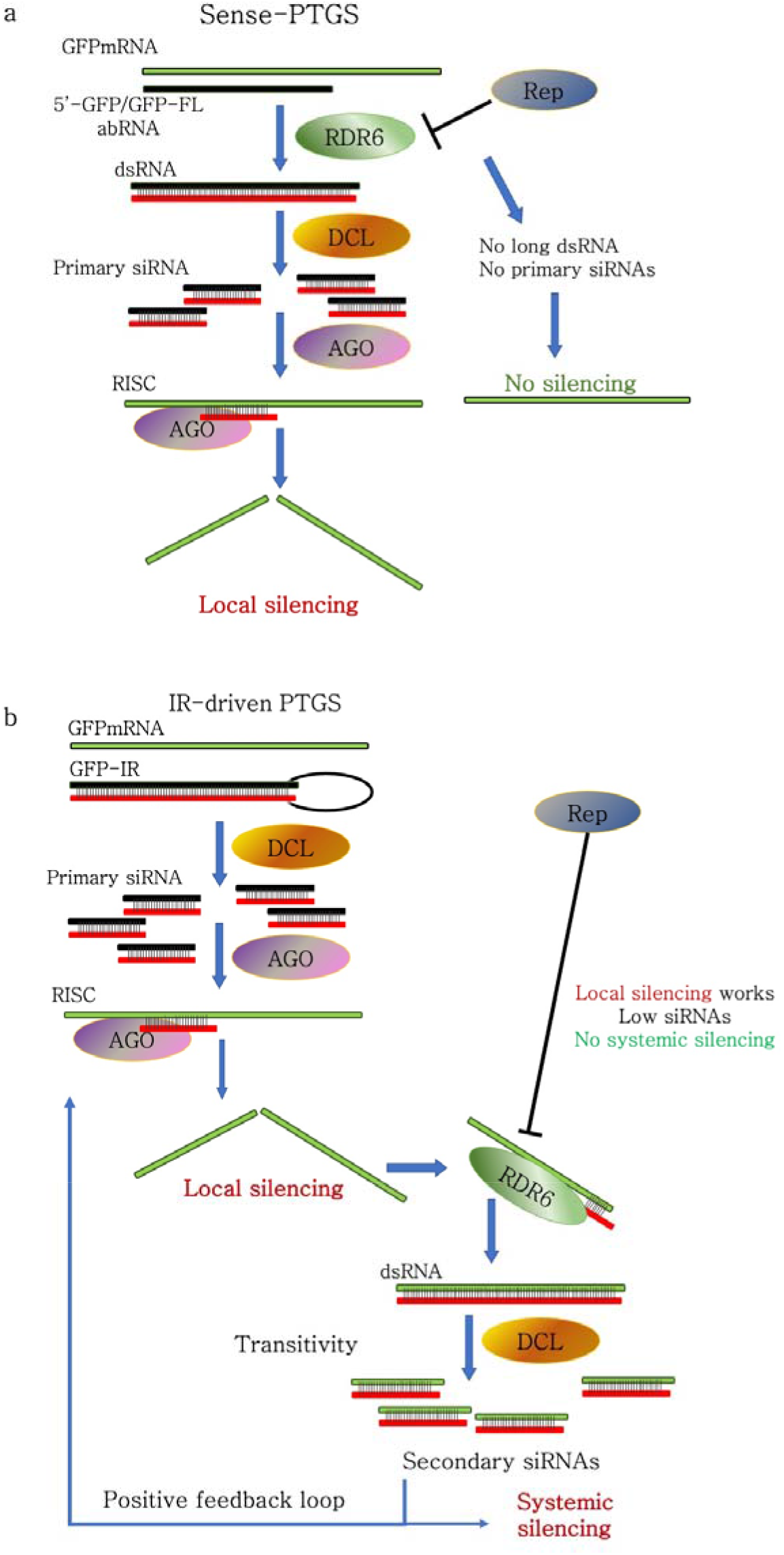
Proposed model for the mode of action of BCTIV-Rep VSR activity. a) S-PTGS is triggered by the RDR6-driven dsRNA synthesis using aberrant RNA (abRNA) as a template. Both 5’-GFP and GFP-FL dsRNAs are cleaved into sRNAs by DCLs. The sRNAs are loaded onto AGO proteins for cleavage of the target GFP mRNA. In the presence of Rep, S-PTGS mechanism does not function. GFP mRNA Northern blot shows that Rep represses the cleavage of GFP mRNA by S-PTGS inducer. Moreover, sRNA Northern Blot showed that Rep also suppresses the production of sRNAs. Therefore, Rep-VSR activity should be associated with the inhibition of DCL or RDR6 activity b) PTGS induced by GFP-IR shows that Rep cannot block the local silencing, triggered by DCL cleave of the GFP-IR, suggesting that Rep does not interfere with the DCL activity, which leaves RDR6 as the only possible target of Rep. Moreover, Rep still reduces the sRNAs produced GFP-IR, suggesting that the propagation of the sRNA production (positive feedback loop) by RDR6 activity is mitigated by Rep, pointing out the interaction between Rep and RDR6.

## DISCUSSION

RNA silencing is a cellular defense mechanism conserved across kingdoms, which is triggered by dsRNA in plants (Zhao *et al*. 2022). In spite of being ssDNA viruses with no dsRNA phase in their replication cycle, geminiviruses have been shown to be targeted by plants via post-transcriptional gene silencing by producing bi-directional transcription from opposite polarity or the strong fold-back structure of transcripts (Vanitharani et al. 2005). In addition, viral gene expression can lead to aberrant transcripts, which are further processed by RDRs to form dsRNAs. Therefore, geminiviruses must confront posttranscriptional gene silencing (PTGS) to achieve successful infections (Luna et al. 2017). Several open reading frames were identified as PTGS suppressors in geminiviruses but still, there is no report of silencing suppressors in BCTIV (Ramesh et al. 2017). In this study, we used distinct approaches to dissect the role of BCTIV genes in the suppression of PTGS.

When the GFP-FL is agroinfiltrated on WT plants, it induces GFP expression, but it also silences itself due to the production of aberrant RNAs, triggering S-PTGS (Parent et al. 2015). When GFP-FL is transiently expressed, we showed that BCTIV-V2 works as an effective silencing suppressor showing overlapping features with TBSV-P19 VSR activity whereas Rep also represses the self-silencing of FL-GFP but only very mildly (Figure 2). However, when the GFP is solely expressed in the genome in 16C *N. benthamiana* (without transient GFP-FL expression) and the S-PTGS is induced by transient expression of 5′-GFP, TBSV-P19 and BCTIV-V2 have no repressive effect on the silencing, whereas BCTIV-Rep substantially suppresses the S-PTGS (Figure 1). The Northern blots of leaves in which S-PGTS was induced by infiltration with 5′-GFP, TBSV-P19, and BCTIV-V2 co-infiltrated samples showed accumulation of sRNAs, whereas BCTIV-Rep co-infiltrated sample was almost fully devoid of sRNAs (Figure 1). The sRNA signature indicates that P19 and V2 do not interfere with the sRNA production substantially. On the other hand, Rep interferes with sRNA production. These observations implicate that BCTIV-Rep has a distinct mechanism of action from BCTIV-V2 and TBSV-P19.

GFP-IR is a very strong silencing inducer, which can lead to systemic silencing within 10 dpi. Upon transient expression of GFP-IR, due to the hairpin structure, dsRNA immediately forms and is subsequently processed by the DCL proteins. In this situation, none of the silencing suppressors, including Rep could repress the local silencing induced by GFP-IR (Figure 3). Nevertheless, the sRNA Northern blot showed that Rep and C1 could decrease the amount of sRNA produced upon GFP-IR expression. Although the decrease did not lead to suppression of local silencing, the systemic silencing phenotype was consequently affected. Contrary to Rep, V2 did not have any effect on systemic silencing. It has recently been proposed that the AGO proteins sequester the sRNAs and mobile sRNAs can induce systemic silencing only when their concentration is above a certain threshold, determined by the concentration of AGO proteins (Vionnet 2022). Therefore, the reduction in sRNA level by Rep which does not affect local silencing is low enough to avoid induction of systemic silencing. Besides, it is worth mentioning that the necrotic phenotype of the Rep and C1 could further decrease the production of sRNAs after 6dpi, which contributes at least partly to the lack of systemic silencing phenotype in these co-infiltrations.

Nevertheless, it was still puzzling that P19 did not lead to any silencing suppression when 5′-GFP or GFP-IR was expressed on stably GFP expressing 16C plants (Supplementary Figure 3a, b). The vast majority of the characterization of the P19 activity was performed under transient expression of GFP in 16C or WT backgrounds (Martin-Hernandez and Baulcombe, 2008). Confirming the previous literature, when GFP was transiently expressed, we could recapitulate the silencing suppression phenotype of p19 (and V2) on WT or 16C plants (Vargason et al. 2003, Silhavy et al. 2002) (Supplementary Figure 3c-f). The underlying mechanism of the VSR activity of TBSV-P19 on S-PTGS depending on the presence of transient GFP-FL expression remains enigmatic (Supplementary Figure 3a-f). In order to understand the phenomenon better, we transiently expressed P19 two and one day before the GFP-IR expression on 16C samples. In three independent experiments, we consistently found out that P19 can act as a silencing suppressor in the absence of GFP-FL, if it is expressed not one day but at least two days before the silencing inducer GFP-IR or 5’-GFP is expressed on 16C plants (Supplementary Fig 4a-e). Although p19 suppresses silencing when co-infiltrated with GFP-FL, when GFP-FL is expressed one or two days before p19, p19 cannot suppress the silencing (Supplementary Figure 4f-i). It has previously been demonstrated that P19 binds to ds-siRNA to inhibit the silencing derived from ds-siRNA (Vargason et al. 2003). Also, in line with P19-binding specificity, P19 failed to inhibit the activation of RISC by ss-siRNA, similarly, P19 was not able to suppress the target cleavage of the already activated RISC (Lakatos et al. 2004). Furthermore, another study showed CymRSV infection was not able to revert the already established GFP RNA silencing. (Silhavy et al. 2002). In summary, our experiments as well as the given literature indicate that the time of p19 expression relative to the GFP-FL is critical for its function. The difference in timing may create a difference in p19 and siRNA concentrations at a given time, which is critical for the VSR activity.

Although this is the first report of silencing suppressor gene of BCTIV-Rep protein, Rep was previously identified as a PTGS suppressor in different geminiviruses like *Wheat dwarf virus* (WDV) from *Mastrevirus* (Kasschau and Carrington 1998, Liu et al. 2014). Yet, BCTIV-Rep protein is only very distantly related to WDV-Rep proteins based on nucleic acid sequence (Supplementary Figure 5). Therefore, the precise identification of the RNAi pathways that Rep proteins involve in may shed light on uncharted functions of Rep proteins in geminiviruses. Besides, Rep proteins from TYLCV, TGMV, and ACMV were shown to interfere with the plant DNA methylation machinery and suppress transcriptional gene silencing (Rodríguez-Negrete et al. 2013). However, the role of BCTIV-Rep protein on TGS remains unexplored. Moreover, V2 and its homolog AV2 have previously been identified as a suppressor in Geminiviruses, with a weaker silencing outcome than P19 (Sharma et al. 2010), but it had not been reported for BCTIV before this study.

To highlight the mode of action of PTGS suppressor activity of BCTIV, we propose that BCTIV-Rep protein interferes with RDR6 (Figure 7). In S-PTGS, where WT and 16C *N*.*benthamiana* inoculated with FL-GFP and 5′GFP, respectively, RDR6 is recruited to produce dsRNA from the aberrant GFP transcripts (Figure 5a). Rep protein suppressed the local silencing derived from RDR6 mediated dsRNA production. Therefore, 5′-GFP infiltrations do not lead to sRNA production when co-infiltrated with Rep. Unlike during S-PTGS, in GFP-IR induced PTGS, the hairpin transcript is already dsRNA, therefore RDR6 is not required for the production of primary siRNAs, which leads to local silencing (Figure 5b) and in consequence BCTIV-Rep does not suppress IR-induced PGTS. It is noteworthy, however, that RDR6 is also indispensable for systemic silencing (Harmoko et al. 2013, Vaistij and Jones 2009, Dalakouras et al. 2020). The model where Rep interferes with RDR6 activity can explain our current observations, but further experiments are needed to prove the direct or indirect interaction of Rep and RDR6.

The Rep protein of WDV has been functionally characterized. Truncations of WDV-Rep at the N terminus (9 amino acids) and the C-terminus (30 amino acids) did not affect the silencing suppression function and the N-terminus was critical for the necrotic phenotype (Wang et al. 2014). On the other hand, we have further characterized BCTIV-Rep with truncating domains on both the N and the C terminus. In the line with previous findings in WDV, the pathogenicity determinant domain of BCTIV-Rep is also located on the N-terminus. However, we found that all truncated Rep proteins we produced, did not maintain the suppression of the silencing phenotype of BCTIV-Rep, except that C-terminus is shown to have a limited effect on the silencing suppression of Rep.

## CONCLUSION

In this study, we determined the BCTIV-Rep and -V2 as viral suppressors of PTGS. We showed that Rep reduces the siRNA accumulation upon transient and stable expression of the target gene. On the other hand, BCTIV-V2 protein also has S-PTGS suppression activity only if GFP-FL is also transiently expressed, which is similar to its homologous geminivirus-V2 proteins and TBSV-P19.

## Data Availability Statement

The coding sequences of the BCTIV genes are provided in the text. In addition, plasmids used in this study are available upon request.

## Author Contributions

OE, GK, MW, and VVU designed the project. SE, AB, ZY conducted experiments. SE, VVU, and OE analyzed the data. VVU, GK, and SE wrote the manuscript. All authors analyzed the data and reviewed the manuscript. VVU supervised the study.

## Acknowledgements

We thank Arvid Hanke and the other Epigenetics Lab members at RLP AgroScience GmbH for the careful revision of the data and the manuscript.

