## Supplementary Figures for "*Beet curly top Iran virus* Rep and V2 gene work as suppressors of post-transcriptional gene silencing through separate mechanisms"

Ebrahimi et al 2022, Supplementary Data

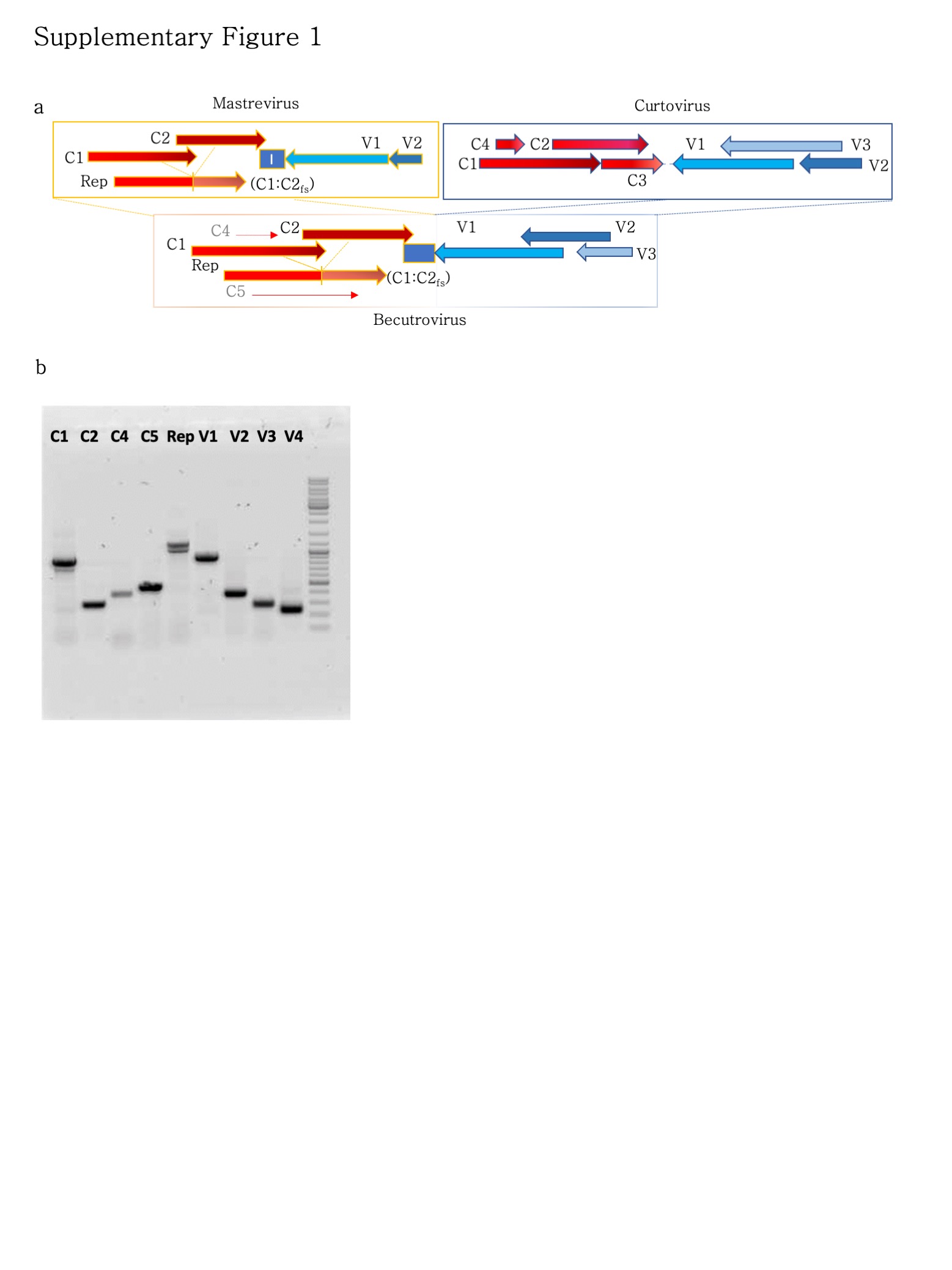

**Supplementary Figure 1| BCTIV genome organization and proof of expression of BCTIV genes upon agroinfiltration.** a) The sketch on the top left shows the genome organization of mastrevirus. The sketch on the top right shows the genome organization of curtovirus. The bottom row highlights the similarity C-strand similarity of becurtovirus to mastrevirus and the V-strand similarity of becurtovirus to curtovirus. b) RT-PCR results show that the BCTIV genes are successfully expressed in the plant after agroinfiltrations. The agroinfiltrated *N.benthamiana* leaf material is collected at 6dpi. RNA extraction, DNase I treatment is applied on the samples, followed by the RNA clean-up. The samples are amplified by one-step SuperScript™ III One-Step RT-PCR System with Platinum™ Taq (Thermofisher). Hypothetical genes, C5 and V4 were not used in this study.

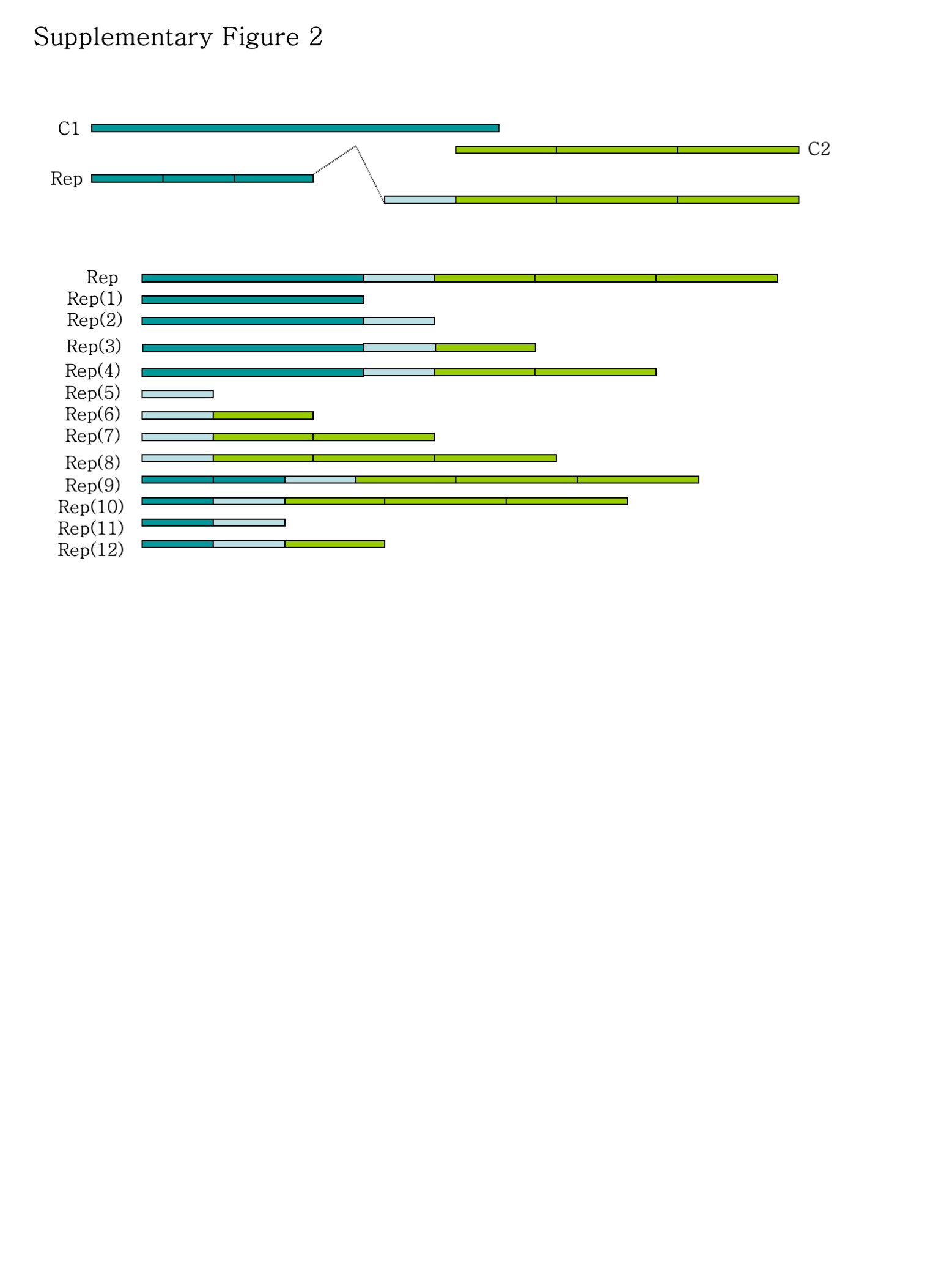

**Supplementary Figure 2| BCTIV-Rep truncations scheme.** Rep gene contains three main expression domains. (1) The domain, which overlaps with the N terminus of C1 protein, is shown in turquoise (2) The new domain, which is a frameshift of C1 mid-region, is shown in light blue. (3) The frameshifted C2 domain, shown in green. Domain (1) and Domain (3) are divided into three subfragments and in total 12 Rep truncations are obtained and functionally analyzed.

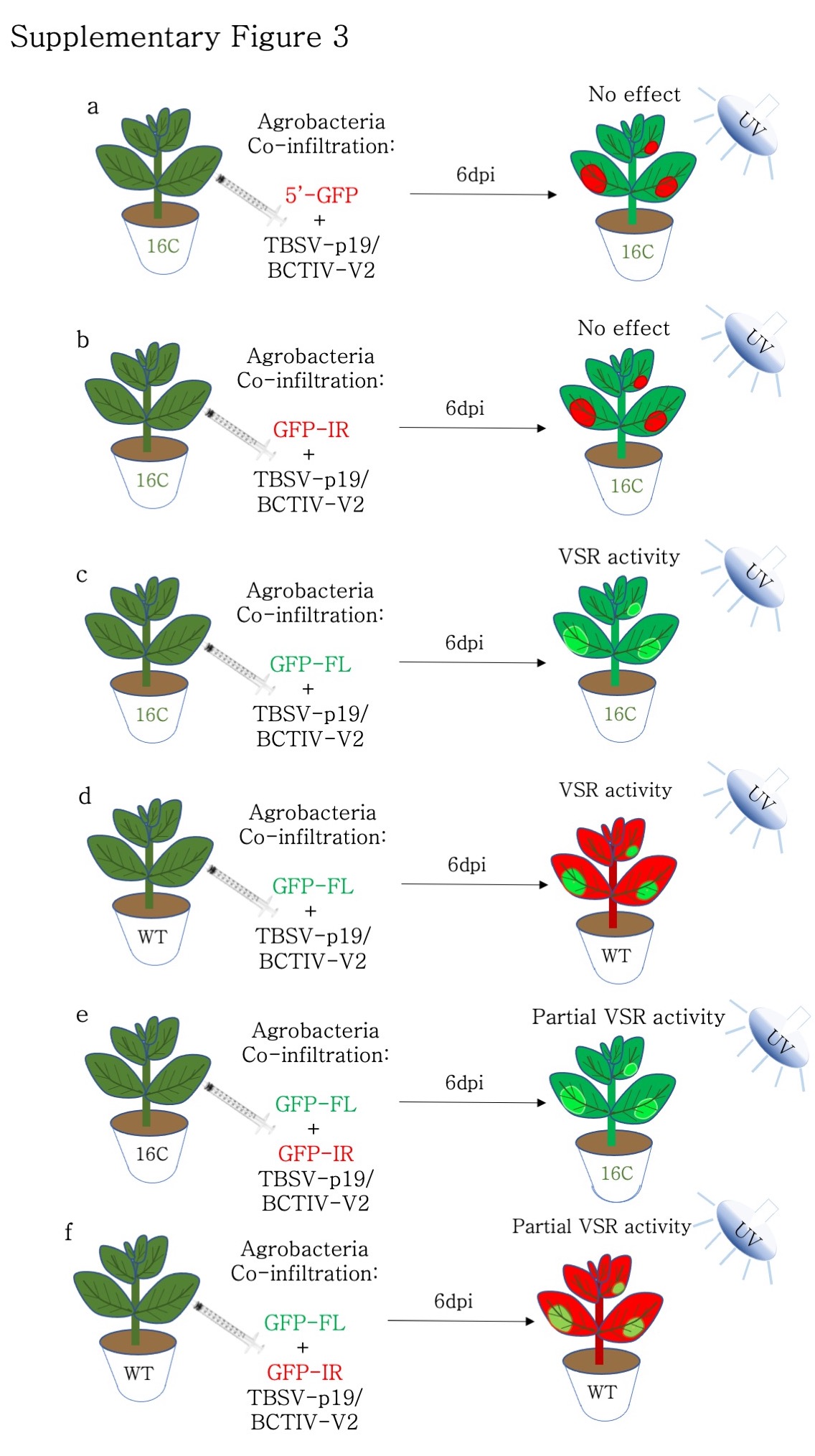

**Supplementary Figure 3| Summary of the VSR activity of TBSV-p19 and BCTIV-V2 screen.** a) When S-PTGS is induced by 5’-GFP on 16C plants, neither p19 nor V2 can suppress S-PTGS. b) When PTGS is induced by GFP-IR on 16C plants, neither p19 nor V2 can suppress S-PTGS. c) Both p19 and V2 can suppress S-PTGS, triggered by GFP-FL on 16C plants d) Both p19 and V2 can suppress S-PTGS, triggered by GFP-FL on WT plants. e) p19 and V2 can partially suppress the GFP-FL (S-PTGS inducer) and GFP-IR (PTGS inducer) on 16C plants. f) p19 and V2 can partially suppress the GFP-FL (S-PTGS inducer) and GFP-IR (PTGS inducer) on WT plants.

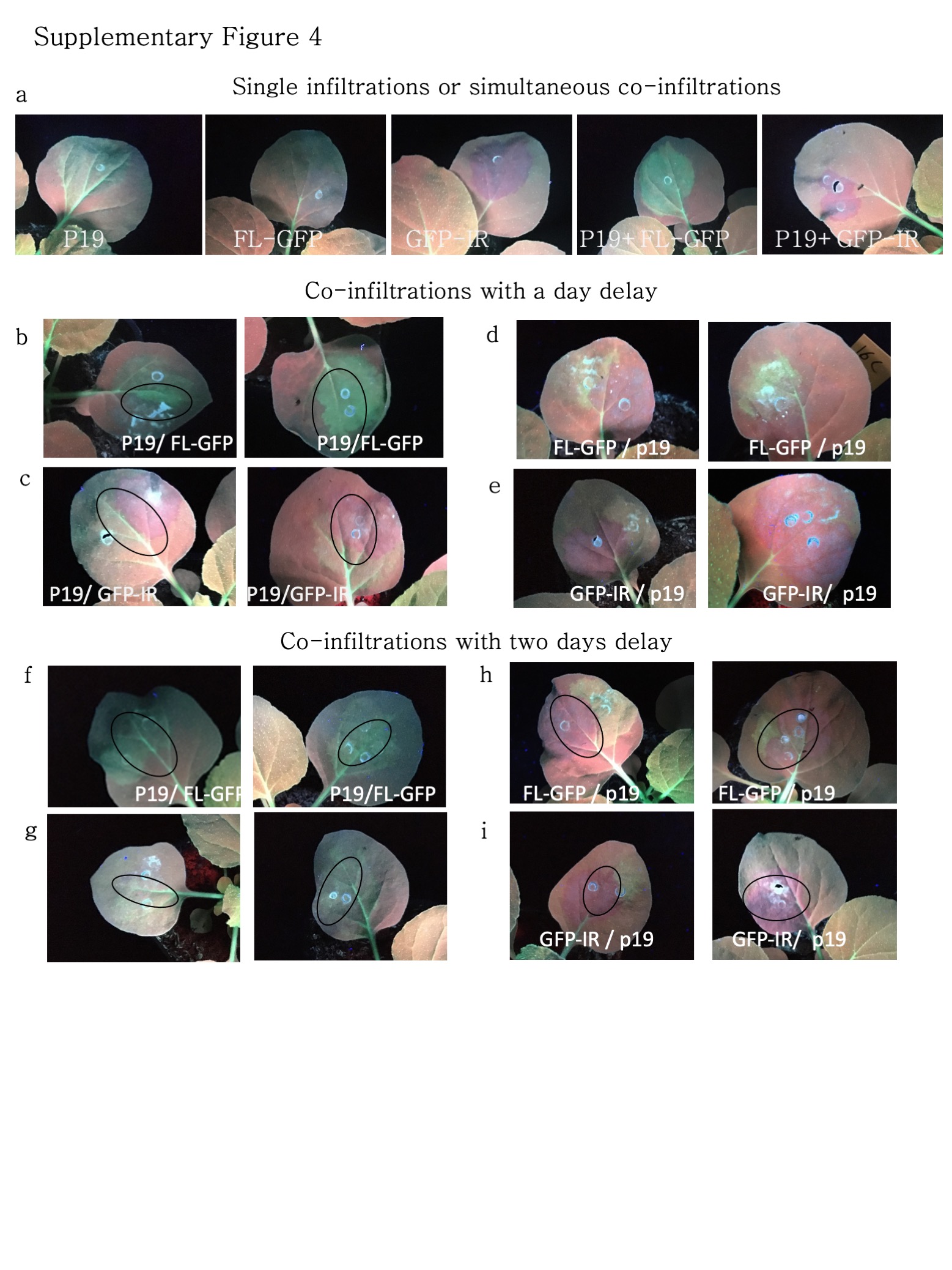

**Supplementary Figure 4| Agroinfiltration scheme for different timing of p19 expression with regards to the silencing inducers** All the plant images are obtained under a UV lamp using a DSLR camera at 6 dpi based on the infiltration of silencing inducers. a) The leaves of the 16C *N.benthamiana* plants, either infiltrated with a single expression vector or simultaneous co-expression of two expression vectors such as p19 and FL-GFP. b) 16C leaves where p19 is infiltrated a day before GFP-FL (S-PTGS inducer) with partially overlapping infiltration domains show silencing suppression. c) 16C leaves where p19 is infiltrated a day before GFP-IR (PTGS inducer) with partially overlapping infiltration domains shows no suppression at the overlapping domains. d) 16C leaves where GFP-FL is infiltrated a day before p19 with partially overlapping infiltration domains show silencing in the overlapping domains e) 16C leaves where GFP-FL is infiltrated a day before p19 with partially overlapping infiltration domains shows no sign of silencing suppression f) 16C leaves where p19 is infiltrated two days before GFP-FL, S-PTGS inducer with partially overlapping infiltration domains shows silencing suppression. g) 16C leaves where p19 is infiltrated two days before GFP-IR (PTGS inducer) with partially overlapping infiltration domains show major suppression of silencing. h) 16C leaves where GFP-FL is infiltrated two days before p19 with partially overlapping infiltration domains show no silencing suppression i) 16C leaves where GFP-FL is infiltrated two days before p19 with partially overlapping infiltration domains also show no sign of silencing suppression.

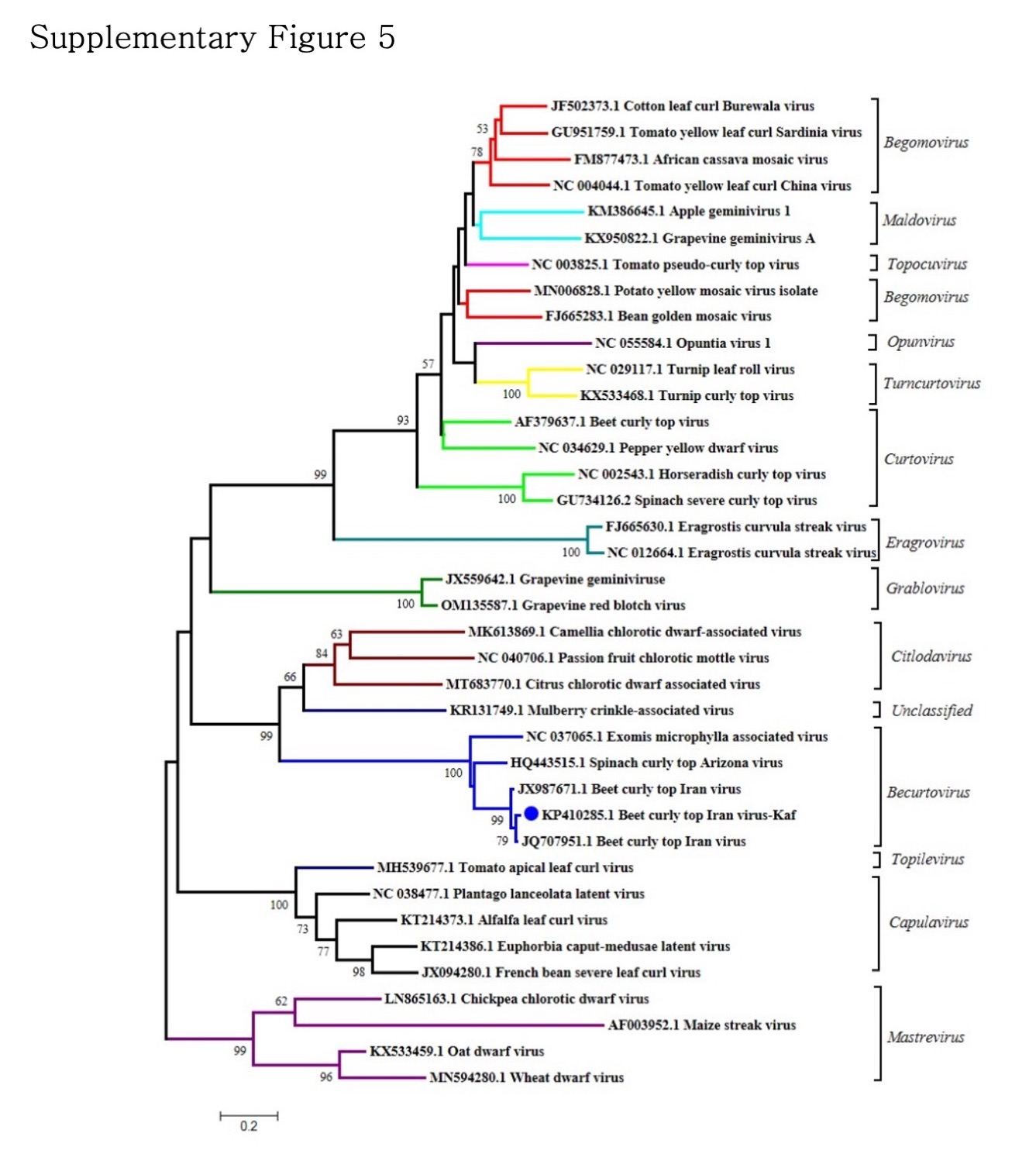

**Supplementary Figure 5|Phylogenetic Tree of Geminivirus Rep proteins.** Phylogenetic tree based on the neighbor-joining method is obtained using the nucleotide sequence of the Rep genes or homologous genes in geminividae.
